# In vivo assay of cortical microcircuitry in frontotemporal dementia: a platform for experimental medicine studies

**DOI:** 10.1101/416388

**Authors:** Alexander D Shaw, Laura E Hughes, Rosalyn Moran, Ian Coyle-Gilchrist, Tim Rittman, James B Rowe

## Abstract

The analysis of neural circuits can provide critical insights into the mechanisms of neurodegeneration and dementias, and offer potential quantitative biological tools to assess novel therapeutics. Here we use behavioural variant frontotemporal dementia (bvFTD) as a model disease. We demonstrate that inversion of canonical microcircuit models to non-invasive human magnetoecphalography can identify the regional- and laminar-specificity of bvFTD pathophysiology, and their parameters can accurately differentiate patients from matched healthy controls. Using such models, we show that changes in local coupling in frontotemporal dementia underlie the failure to adequately establish sensory predictions, leading to altered prediction error responses in a cortical information-processing hierarchy. Using machine learning, this model-based approach provided greater case-control classification accuracy than conventional evoked cortical responses. We suggest that this approach provides an *in vivo* platform for testing mechanistic hypotheses about disease progression and pharmacotherapeutics.

## Introduction

The impairment of brain circuit physiology occurs early in neurodegeneration. For example, the loss of synapses, synaptic plasticity, and effective information processing in microcircuits precede the onset of atrophy and behavioural change in animal models of neurodegeneration (Rowan et al. 2003; Hof and Morrison 2004). New quantitative tools to assay these early changes are a key goal for the development and monitoring of therapies to slow or prevent neurodegenerative disease.

There is strong preclinical evidence of functional impairment in neural circuits before cell death or atrophy, including the downstream effects of oligomeric modified and misfolded proteins on axonal transport, synapse density and plasticity (Wilcock et al. 2009; Castillo-Carranza et al. 2015). In humans however, the equivalent physiological observations have been limited by the low resolution and indirect nature of brain imaging, such as structural and functional magnetic resonance imaging (MRI) (De Jong et al. 2008) and evoked responses in electroencephalography (EEG) or magnetoencephalography (MEG) (Stam 2005, 2010; Hughes and Rowe 2013). Nonetheless, there is growing evidence for the reorganisation of brain networks, and change in the efficiency of information processing, in patients with Alzheimer’s disease (Zhou et al. 2010; Sami et al. 2018), Parkinson’s disease (Crossley et al. 2014), progressive supranucelar palsy (Rittman et al. 2016; Cope et al. 2018) and frontotemporal dementia (Hughes et al. 2013, 2018).

Recent advances in computational models of human neural circuits offer new tools for *in vivo* assays of cortical function, with increasingly detailed anatomical and pharmacological specificity (Moran, Jung, et al. 2011; Moran, Symmonds, et al. 2011; Bastos et al. 2012). Neurophysiologically informed modelling goes beyond descriptive biomarkers by providing a mechanistic link to realistic microscopic processes embedded within the model. For example, the canonical microcircuit model (CMC) of cortical columns comprises layer-specific and inter-connected populations of pyramidal cells, stellate cells and inhibitory interneurons (Douglas and Martin 1991; Haeusler and Maass 2007), which link the dynamics of macroscopic brain activity to network parameters describing the interactions amongst subpopulations. In both human and animal brain imaging, it has been shown that the CMC model accurately recapitulates mechanisms known to be interrupted by distinct genetic (Gilbert et al. 2016) and disease (Hughes and Rowe 2013; Cooray et al. 2015; Symmonds et al. 2018) loci. Moreover the model has been validated pharmacologically using modulators of AMPA, GABA and NMDA receptors to demonstrate veridical parameter recovery (Sc et al. 2010; Moran, Jung, et al. 2011; Moran et al. 2014; Muthukumaraswamy et al. 2015).

The inversion of such CMC models, constrained by empirical brain imaging data, has significant advantages over historical approaches to evoked and induced studies applied typically in the context of EEG and MEG. Evoked responses and spectral densities are limited in the biological information that they yield and lack the biological detail required to test mechanistic questions about disease or treatment. However this difference in feature space suggests that mechanisms must differ at a neuronal level. The outlined modelling approach takes advantage of this and so in contrast to data feature reporting, biological models such as the CMC attempt to explain differences in evoked responses or spectra giving insight from neurophysiological data in terms of the parameterised and biologically plausible circuits that can generate the observed invasive (LFP), scalp (EEG) or sensor (MEG) data (e.g. 19).

We applied this modelling approach to examine neurodegenerative disease, using the behavioural variant of frontotemporal dementia (bvFTD) as a demonstrator condition. We selected bvFTD as a human disease model because of its regional and laminar specificity within the cortex. Behavioural variant frontotemporal dementia (bvFTD) is a severe neurodegenerative disorder characterised by progressive deterioration of behaviour and personality (Bang et al. 2015), with heterogeneous molecular pathology involving misfolding and aggregation of either TAR DNA-binding protein 43 (TDP-43), microtubule associated protein Tau, or rarely fused-in-sarcoma protein (Neary et al. 2005). In addition, preclinical models demonstrate common downstream consequences including changes in synaptic morphology, signalling and density and cell death. Interestingly, in humans and transgenic models, cell death in frontal and temporal regions is most marked in superficial cortical layers (II-III) (Kersaitis et al. 2004), as well as in layer V in selective frontal regions (Kim et al. 2012; Santillo and Englund 2014), providing clear testable hypotheses for the inversion of CMC models.

To probe neural circuits in bvFTD, we studied patients during a passive auditory oddball paradigm. Auditory stimuli were either standard tones, or which deviated in one of five dimensions (frequency, loudness, laterality, duration, or a central silent period). Evoked responses to deviant tones, and large-scale cortical interactions (Hughes and Rowe 2013) during such auditory oddball paradigms are grossly abnormal in bvFTD and related disorders. There is an extensive literature on the effects of neurological and psychiatric (Umbricht and Krljesb 2005) disease and ageing (Naatanen et al. 2011) on the ‘mismatch negativity response (MMN)’, to deviant vs. standard tones. The neural generators of the MMN have been successfully modelled in humans (Garrido et al. 2009; Hughes et al. 2013; Phillips et al. 2015) and validated against invasive electro-corticography (ECog) (Phillips et al. 2016). These biophysically informed models consistently identify a bilateral network of generators including inferior frontal gyrus (IFG), superior temporal gyrus (STG) and primary auditory cortex (A1). In this architecture of the MMN network, the parameters of a biologically informed CMC model include the connection strengths, time constants and cell type contributions to the signal in specific regions and layers of cortex.

Previous studies have confirmed that pateints with bvFTD can undertake this paradigm (Hughes et al. 2013). We applied CMC models to MEG data, in a family of nested neuroanatomical models, using Dynamic Causal Modelling for evoked responses (Friston et al. 2003; Kiebel et al. 2008; Chen et al. 2012). We used the model-evidences, with Bayesian model selection, to identify the most likely model under conventional experimental conditions (standard and deviant tones).

Given an optimised model architecture, we predicted that the model parameters would differ between groups, in accordance with the known laminar- and regional-specificity of bvFTD. Specifically, we tested the hypotheses that (i) the contributions of layers II and V to the evoked response, but not layer IV, are reduced by bvFTD, and (ii) the parameters of connectivity within the regional CMC’s, including the gain of superficial pyramidal cells, accurately distinguish patients from controls.

## Methods

### Participants

We recruited 44 patients with bvFTD meeting consensus diagnostic criteria (Rascovsky et al. 2011) from the Cambridge Centre for Frontotemporal Dementia and Related Disorders. Forty-four healthy controls were recruited from the Medical Research Council Cognition and Brain Sciences Unit volunteer panel. We then subsampled the best age- and sex-matched groups, of 40 per group. The study was approved by the local Research Ethics Committee and all participants gave written informed consent before participation according to the 1991 Declaration of Helsinki.

### Cognitive Examination

All bvFTD patients completed the Addenbrookes Cognitive Examination (Revised) (ACE-R) (Mioshi et al. 2006), which includes subscores for attention, memory, fluency, language and visuo-spatial ability; and the Mini Mental State Examination (MMSE). Patients were further characterised using the Cambridge Behavioural Inventory (CBI), a carer-based questionnaire developed for quantifying the symptom costellation and severity in FTD (Wear et al. 2008).

### MEG Paradigm

Participants were tested on one session each, using a multiple deviant auditory mismatch negativity paradigm (Pakarinen et al. 2004; Hughes et al. 2013). Standard compound sinusoid tones lasted 75 ms duration, of 500, 1000 and 1500 Hz. Deviants differed in either frequency (550, 1100, 1650 Hz), intensity (+/-6dB), duration (25 ms), laterality (missing left or right) or the middle 25ms was omitted (silent gap). Tone-onset-asynchrony was 500 ms. Three bocks of 5 minutes presented a total of 900 standard and 900 deviant trial types.

### MEG pre-processing

All MEG data were collected using a 306-channel Vectorview system (Elekta NeuroMag, Helsinki, Finland) at the MRC Cognitiveion and Brain Sciences Unit with 102 magnetometers, each coupled with 2 planar gradiometers. Data were sampled at 1 KHz and downsampled offline to 500 Hz. Signal separation was achieved using the standardised MaxFilter 2 algorithm (version 2.0, Elekta-Neuromag) prior to conversion to SPM12. Three anatomical fiducial points (the naison and bilateral pre-auricular points) were used for manual coregistration to a T1-weighted magnetic resonance image (individual where available, otherwise SPM template) for source localisation. Five head-position indicator coils and ∼80 head points were generated using a 3D digitiser (Fastrak Polhemus Inc.). SPM was used for artifact rejection with thresholds of 2500 fT and 900 fT for magnetometers and gradiometers, respectively.

Data were epoched -100 to 300 ms around tone onset. Using SPM12, data were band pass filtered 1-40 Hz and a subtracted baseline applied to each trial (−100 to 0 ms). Source localisation was achieved using Smooth priors, a minimum norm solution that uses a smooth source covariance matrix with correlated adjacent sources. From the resultant images, time series were extracted from the 6 locations of interest using previously reported MNI coordinates (Garrido et al. 2008; Phillips et al. 2015): bilateral auditory cortex (MNI coordinates: [-42, -22, 7], [46, -14, 8]), bilateral STG (MNI: [-61, -32, 8], [59, -25, 8]) and bilateral IFG (MNI: [-46, 20, 8], [46, 20, 8]). We used these coordinates in the following way: for each individual, given their own individual source estimates, the local peak of source activity was identified within a 2 mm trap radius around these coordinates in template space. From here the 6 resultant time series were extracted to a pseudo LFP format SPM data file for subsequent DCM analysis by applying the inverse leadfiled. For standard ERP-based analysis of the MMN, average deviant and standard trials were created for each individual and peak amplitude and latency measures for the difference wave (the mismatch response) were extracted between 80 and 200 ms.

### Neural model and connectivity analysis

Dynamic Causal Modelling (DCM) for evoked responses (Kiebel et al. 2008) was employed (SPM12, DCM10) utilising canonical microcircuit models (CMCs) (Douglas and Martin 1991; Bastos et al. 2012) as generative models for each of the 6 regions. The DCM framework permits inversion of a model of data generation, coupling a generative model (*f*) and forward (or spatial, observation) model (*g*):

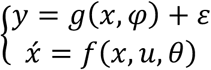

The canonical microcircuit is a special case of convolution-based, mean-field neural mass model (Jansen and Rit 1995; David et al. 2005), comprising four neural populations (superficial layer pyramidal cells, *SP*; granular layer stellate cells, *SS*; deep layer pyramidal cells, *DP*; and inhibitory interneurons, *II*). Each of these populations is described in terms of it’s membrane voltage (*x*_*v*_) and current (*x*_*i*_), governed by sets of parameterised, multivariate first-order differential equations of the form:

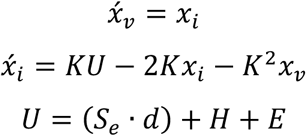

where *K* is the rate-constant of the population, *S*_*e*_ is the extrinsic projections(s) to this layer, *d* = presynaptic firing (calculated using sigmoid activation function with mean field assumption that average *input* is distribution of membrane depolarization over the ensemble), *H* = the sum of postsynaptic-currents targeting this population (i.e. coupling with other populations within this CMC) and *E* = any external / exogenous inputs. The local coupling (G) parameters are depicted in figure 1, while the layer-specific equations of motion are in SupMat1.

**Figure 1.**
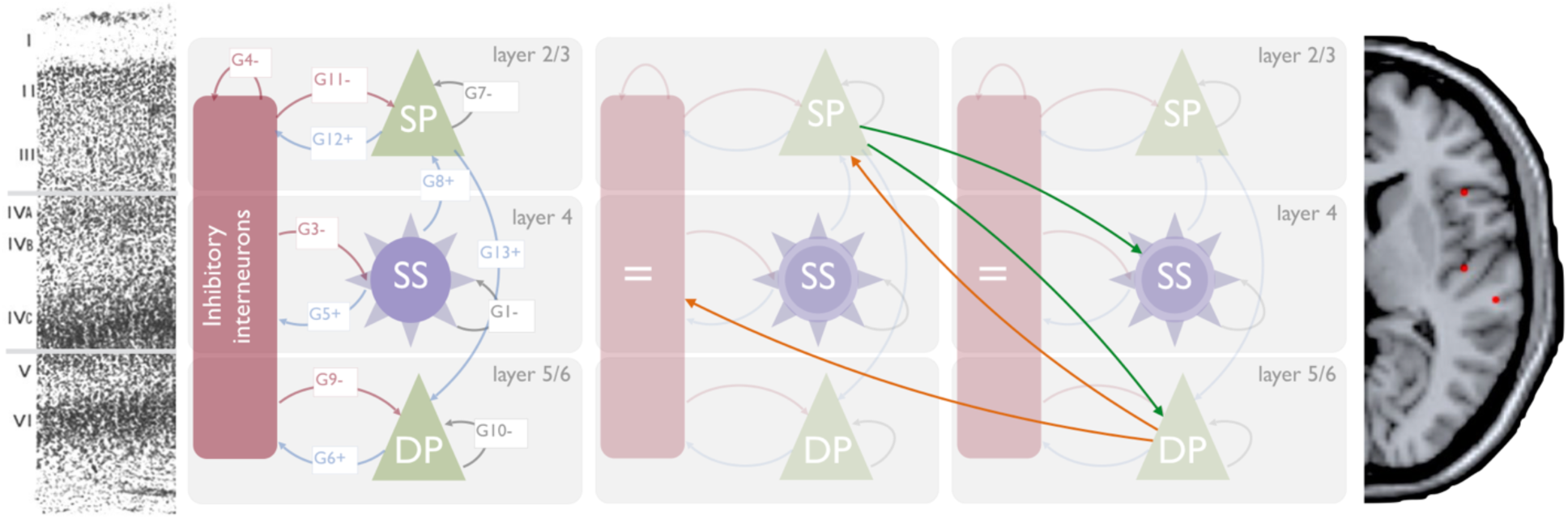
Left: The canonical microcircuit with excitatory (*green*) and inhibitory (*red*) cell populations including pyramidal (*triangle*) and smooth / stellate (*round*) cell types. Blue and red arrows depict intrinsic excitatory and inhibitory connections, respectively. Middle left: histologic depiction of prefrontal cortex cytoarchtecture. Middle Right: Two microcircuits showing extrinsic, layer-specific forward (green) and backward (orange) connections. Right: Template MRI image with red dots marking MNI coordinates for [right] IFG (top), A1 and STG (bottom).

The local-field potential (LFP) forward model was used. This model comprises two parameters: an electrode-gain (*L*) for each CMC (‘node’) in the model and a vector of contribution weights (*J*) for each element of the model state vector, x, such that the full model prediction, *y*, is given by:

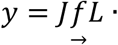

Priors on the contribution weights (*J*) were taken from the literature where only 3 weights were non-zero (and therefore contribute to the signal): SP_V_=0.8, SS_V_=0.2 and DP_V_=0.2. The mapping function, 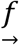, denotes that, in the present model, the Kronecker matrix *J*⨂*L* is mapped to a matrix the size of x before multiplication. This occurs because we enforce similarity across regions in terms of the contributing states (e.g. L2/3 of left and right IFG share the same contribution [*J*] value as A1 (right and left) and STG right and left) and thus the matrix sizes are mismatched. Otherwise the model is as described in Shaw *et al*. (Shaw et al. 2017).

Following Phillips *et al*. (Phillips et al. 2015), 21 plausible model architectures were compared (**figure 2**). These models comprise forward, backward and lateral connections between each of the 6 CMCs. Forward projections originate from SP and target both DP and SS of the target regions whereas backward projections originate in DP and target both SP and II (Bastos et al. 2012; Shipp et al. 2013) (summarised figure 1b).

**Figure 2.**
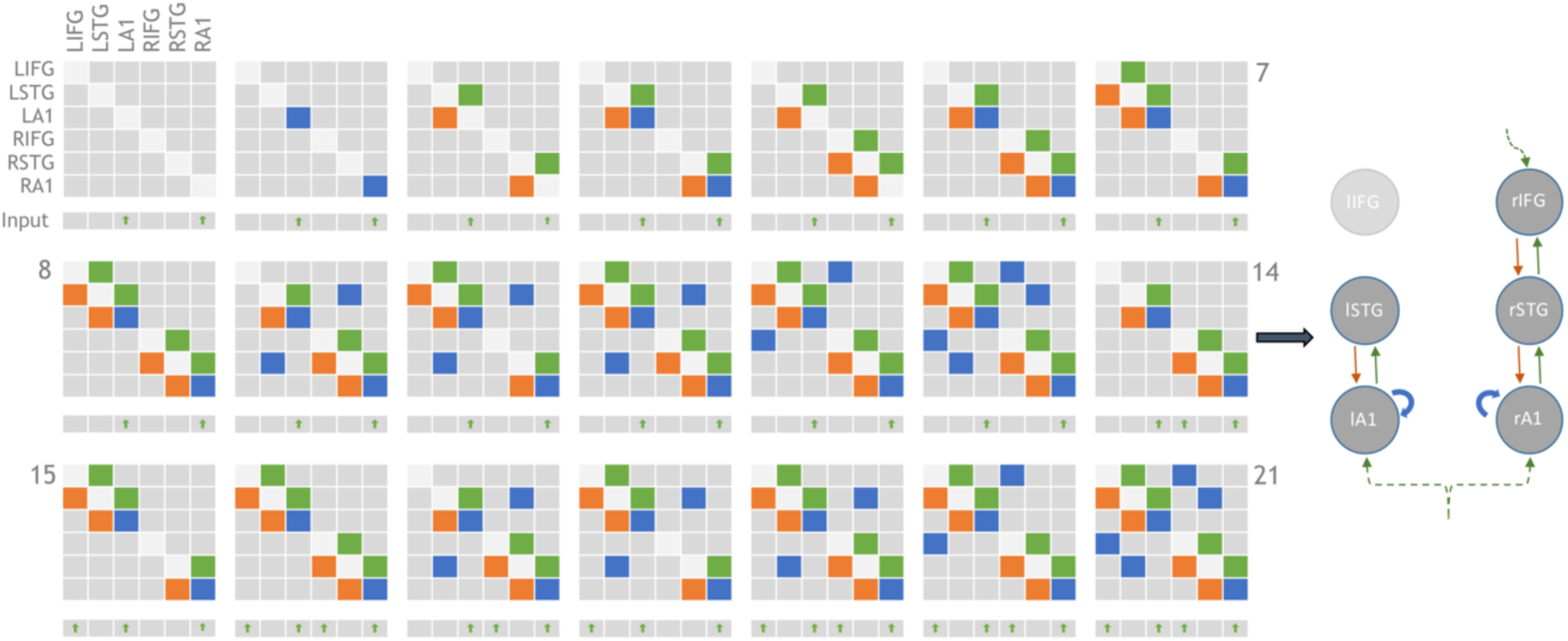
Adjacency matrices showing the 21 model architectures tested, as per Phillips *et al*. 2015. Green, orange and blue blocks represent the presence of forward, backward and lateral (or self) connections modulating the mismatch effect, respectively. L/R-IFG = Left/Right-Inferior frontal gyrus. Inputs are exogenous for sensory regions or endogenous for non-sensory regions. All driving inputs arrive in layer 4 of target regions. Model 14 (depicted right) was the overall winning model, in line with the results of Phillips et al (2015).

The posterior model parameters were estimated by inverting a parameterised full model (generative + forward model). This inversion method is referred to as variational bayes (Friston et al. 2003, 2007), which modulates the log-scaling parameters around static priors (Supplementary table 1).

### SVM pipeline

Support vector machines (LIBSVM implemented in Python (Chang and Lin 2013)) were trained and tested using a permutation-based leave-one-out with replacement cross validation approach. The case excluded for each iteration was selected using the Matlab random integer generator. The SVM was trained and tested on 3 sets of data: 1) the generative model parameters in the form of effective connectivity strengths between nodes (c.f. generative embedding, see (Brodersen et al. 2011)). 2) The forward model parameters in the form of layer-by-node specific population contributions and 3) the amplitudes of the MMN at each region.

## Results

Groups were matched by age (control mean 61.7 range: 45-75; bvFTD mean 60.7 range: 42-78; n.s.) and sex (controls M:F 20:20; bvFTD M:F 21;19; n.s.). Patients were cognitively impaired with average MMSE=23.5/30 (SE 1.0) and ACER-total=69.5/100 (SE 2.9), with typical deficits including severe non-fluency (mean 4.6/14), and milder deficits in attention (mean 14.6/18), memory (mean 15.2/26), language (mean 21.1/24) and visuospatial function (13.3/16) (Figure 3). Contemporay CBI scores were available for 29 patients, with a mean of 85 (+/-50). These scores are qualitatively similar to those of the bvFTD cohort reported by Wear et al (Wear et al. 2008), and are higher than typical CBI scores in Parkinson’s disease, Huntington’s disease and Alzheimer’s disease. Two subjects were excluded retrospectively due to a change of diagnosis while 5 were excluded due to medication changes close to the time of scanning. This resulted in 33 patient datasets and 40 healthy control datasets taken forward for the principal analyses.

**Figure 3.**
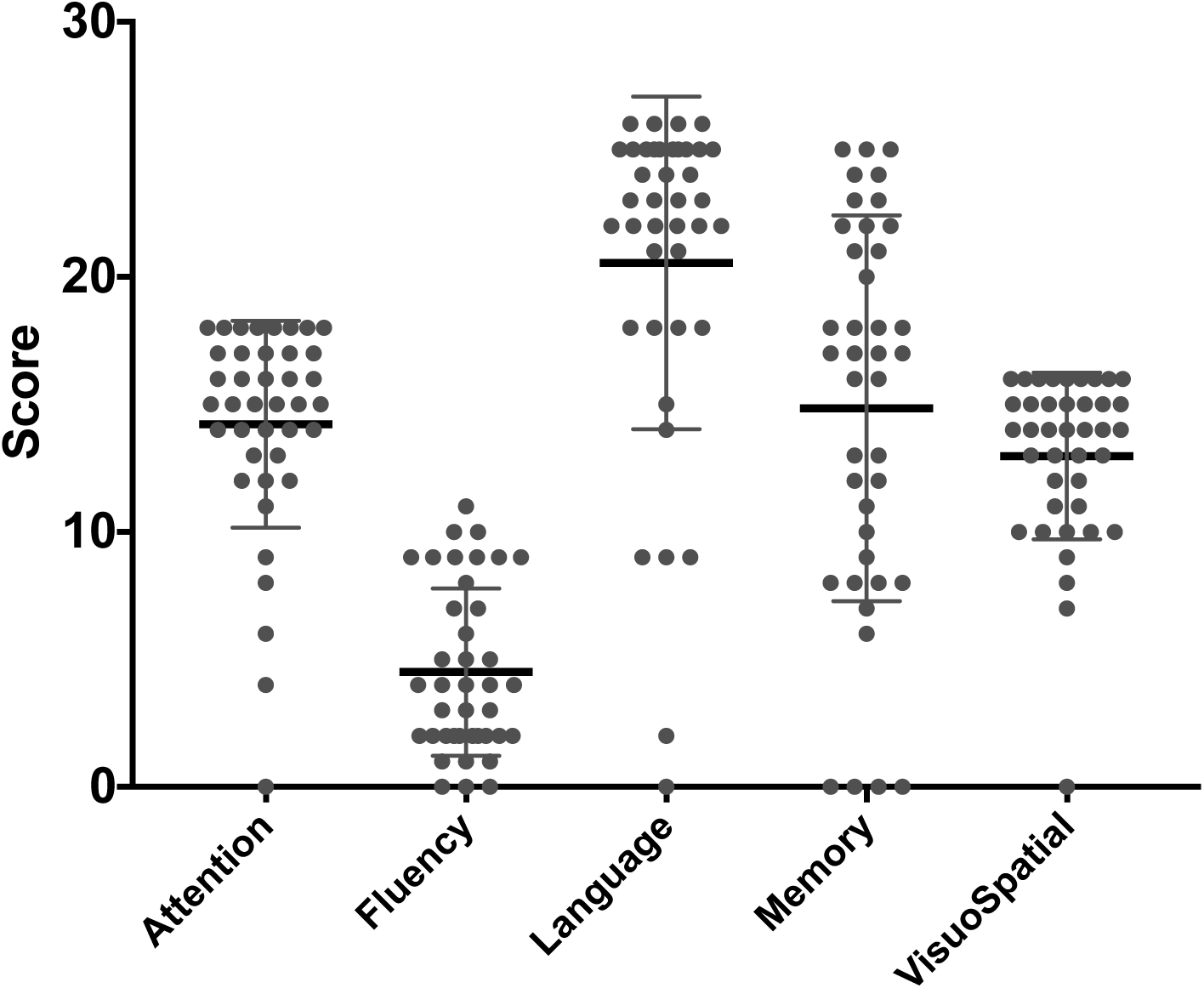
Violin plots of the clinical features from the subsections of ACE-R cognitive examination for the FTD group. Maximum scores are attention, 18, fluency, 14, language, 26, Memory, 26, visuospatial, 16.

Since scanning, at least 15 individuals from the patient cohort have died. Five of these underwent confirmatory post-mortem pathological testing, revealing four cases with TDP43 pathology and one FTLD-tau pathology. In addition, three underwent genetic testing, confirming two with likely TDP43 pathology (C9orf72 hexanucleatide expansions) and one with likely Tau pathology (MAPT mutation).

We confirmed the effect of bvFTD on the MMN event related field, first by averaging over the 6 sources’ timecourse (bilateral IFG, STG and A1) between 80 and 300 ms. A group by condition (2 x 2) analysis of variance (ANOVA) revealed a significant interaction effect for amplitude (F = 9.47, p = 0.002) but not latency (figure 4). Post hoc tests demonstrated that the bvFTD group did not establish an amplitude difference between standard and deviant stimuli (i.e. *the mismatch*) (p=>.05) whereas the control group did (t=-6.2, p<.001).

**Figure 4.**
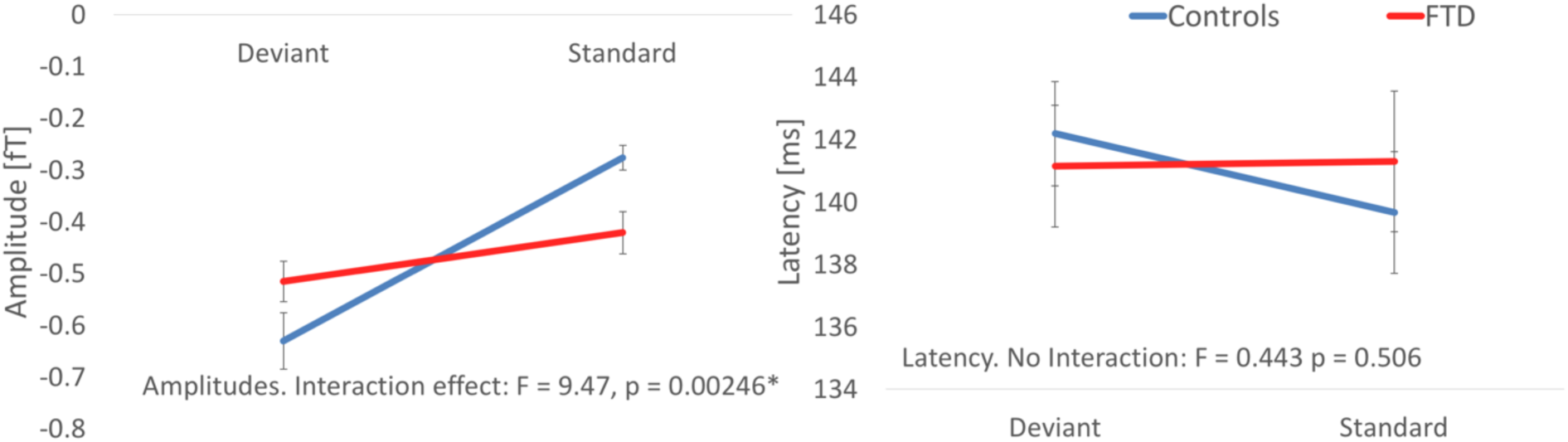
Group changes in amplitude (left) and latency (right) for each condition, averaged over IFG, STG and A1 bilaterally.

Following inversion of the 21 models in figure 2, group data were pooled for Bayesian Model Selection (BMS). BMS was run both with fixed effects and random effects, using a hierarchical family-wise approach. The 21 models were split at three levels (figure 5a), with comparisons performed at each level (RFX and FFX), comprising:

1. Step 1. Models with or without LIFG connectivity (models 7, 8, 10:13, 15, 16, 18:21 vs. 1:6, 9, 14, 17). The family of models without LIFG won in both RFX and FFX analysis (exceedance probability [EP] = 0.89).
2. Step 2. Within the model set without LIFG connectivity, we compared models with or without interhemispheric connections (9,17 vs. 1:6, 14). The family without interhemispheric connections won in both RFX and FFX analysis (EP = 0.68).
3. Step 3. Within the remaining model set, we compared models with or without a top-down (latent) input to rIFG (14 vs. 1:6). The family (model 14 only) with rIFG inputs won in both RFX and FFX analysis (EP = 0.81).

Bayesian model selection was repeated for all subjects (pooled control and FTD groups) over the 21 models (i.e. not family wise). This also converged on model 14, but to test the robustness of this lead model, we undertook 1000 permutations of Bayesian model selection using leave-one-out with replacement cross validation. Model 14 was the lead model 88% of the time followed by model 6, which is nested within model 14 (figure 5b). Model 14 was therefore taken forward for parameter analysis.

**Figure 5a.**
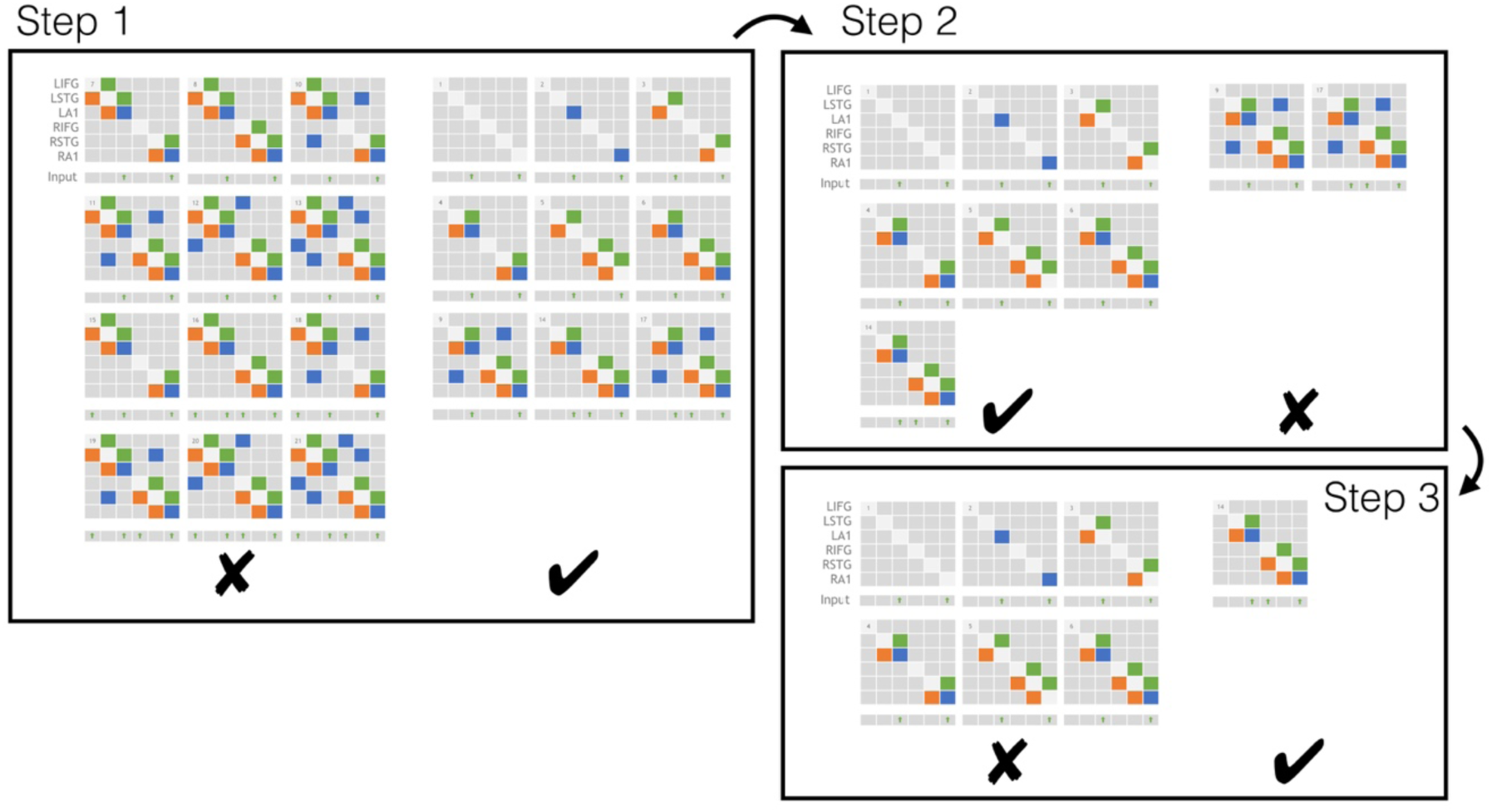
Hierarchical families tested using BMS. Step 1: Models without left IFG perform better than those with (both fixed and random FX). Step 2: Of these models, those without lateral connections perform better than those with (both fixed and random FX). Step 3: Of the remaining 7 models, the model with top-down input performed better than those without (both fixed and random FX).

**Figure 5b.**
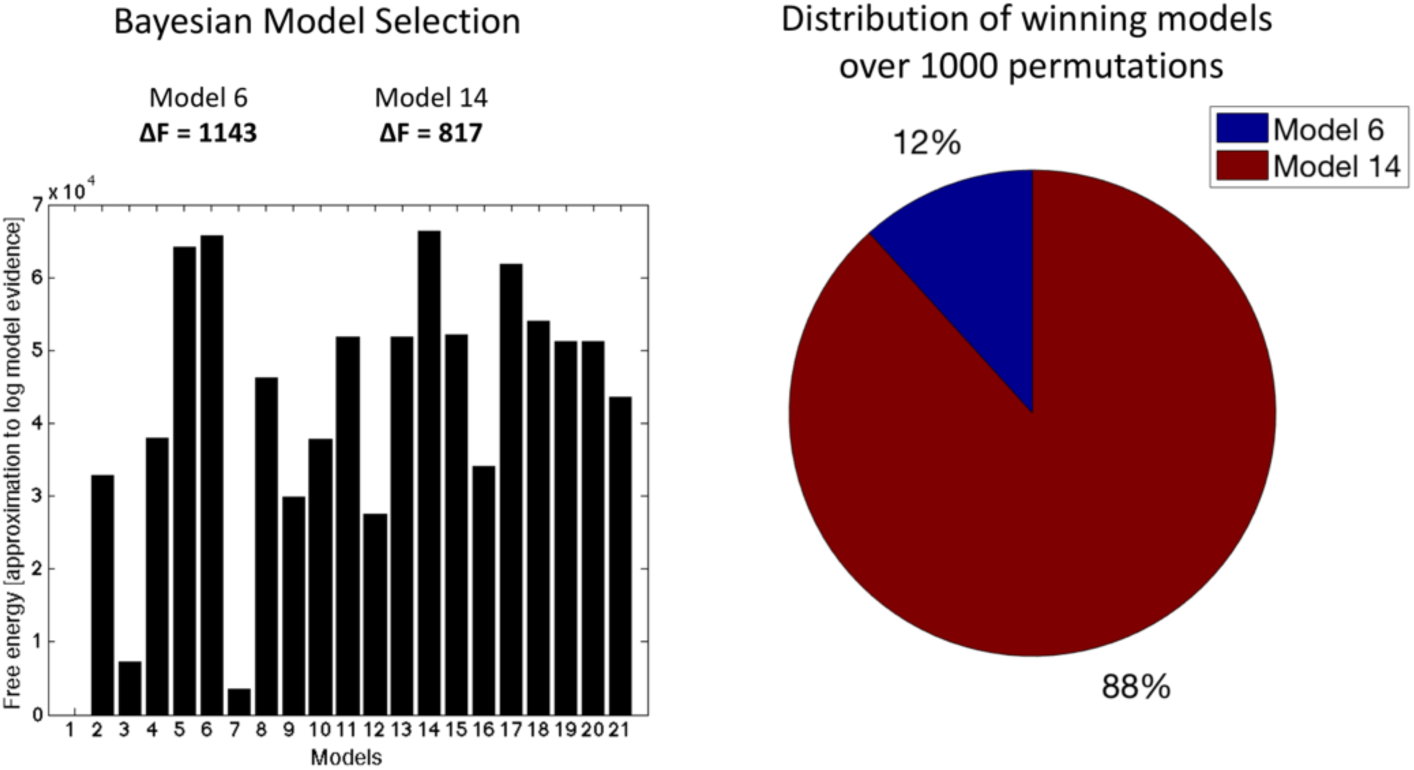
Bayesian model selection for all subjects (pooled groups) over the 21 models (i.e. not family wise) also converges on model 14. One thousand permutations with leave-one-out with replacement were computed with model 14 winning 88% of the time (883 times) followed by model 6, 12% of the time (117 times).

Having identified model 14 as the most likely model architecture, 2 further questions were addressed using the parameters from this model. First, we address the ability of the parameters controlling cortical layer-specific contributions to the MEG signal (‘J’) to differentiate between groups, given the known degenerative pathology in bvFTD (analysis A), based on the evidence of laminar specificity of cell loss in bvFTD. Second, having optimised these ‘contribution’ parameters for each subject, we reinvert the model to estimate local, ‘intrinsic’ coupling strengths between cell populations (analysis B).

### Analysis A: Layer-by-node contributions

Layer-by-node contributions were analysed by ANOVA, which demonstrated a trend towards a group-by-layer interaction (F = 2.6, p = 0.071). Post-hoc independent t-tests revealed a significant reduction of L5/6 STG contribution to the LFP (t=2.8, p=0.005). The parameters did not correlate with ERF amplitudes for either group. No differences were found in the effective connectivity strengths between nodes between groups.

Although the ANOVA of individual layer-by-node contributions did not indicate a strong group difference, these values when taken as a set for classificaiton did separate the groups. Overall classification accuracy (true positive + true negative, table 1) was 99.6% using the layer-by-node population outputs (figure 7). In contrast, generative embedding, using effective connectivity strength between nodes, achieved only 60.7% accuracy, while classification by MMN amplitude was 59.8% accurate (versus 50% by chance).

**Table 1.**
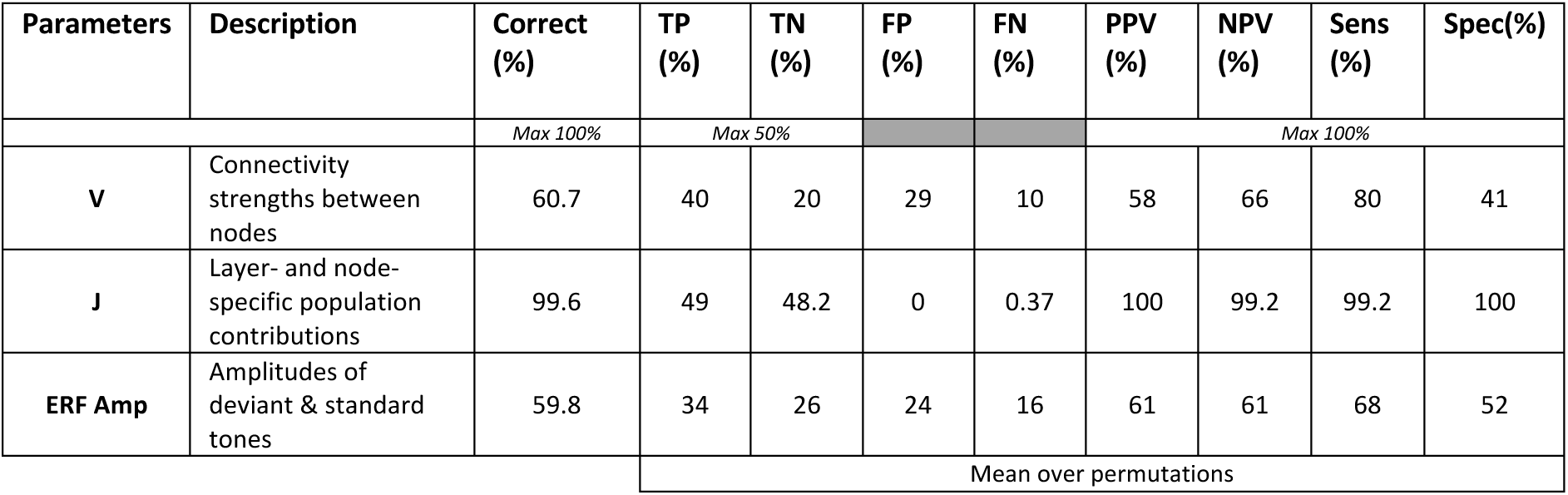
Accuracies (%) and predictive values for the SVM performance across the 3 data.

**Figure 6.**
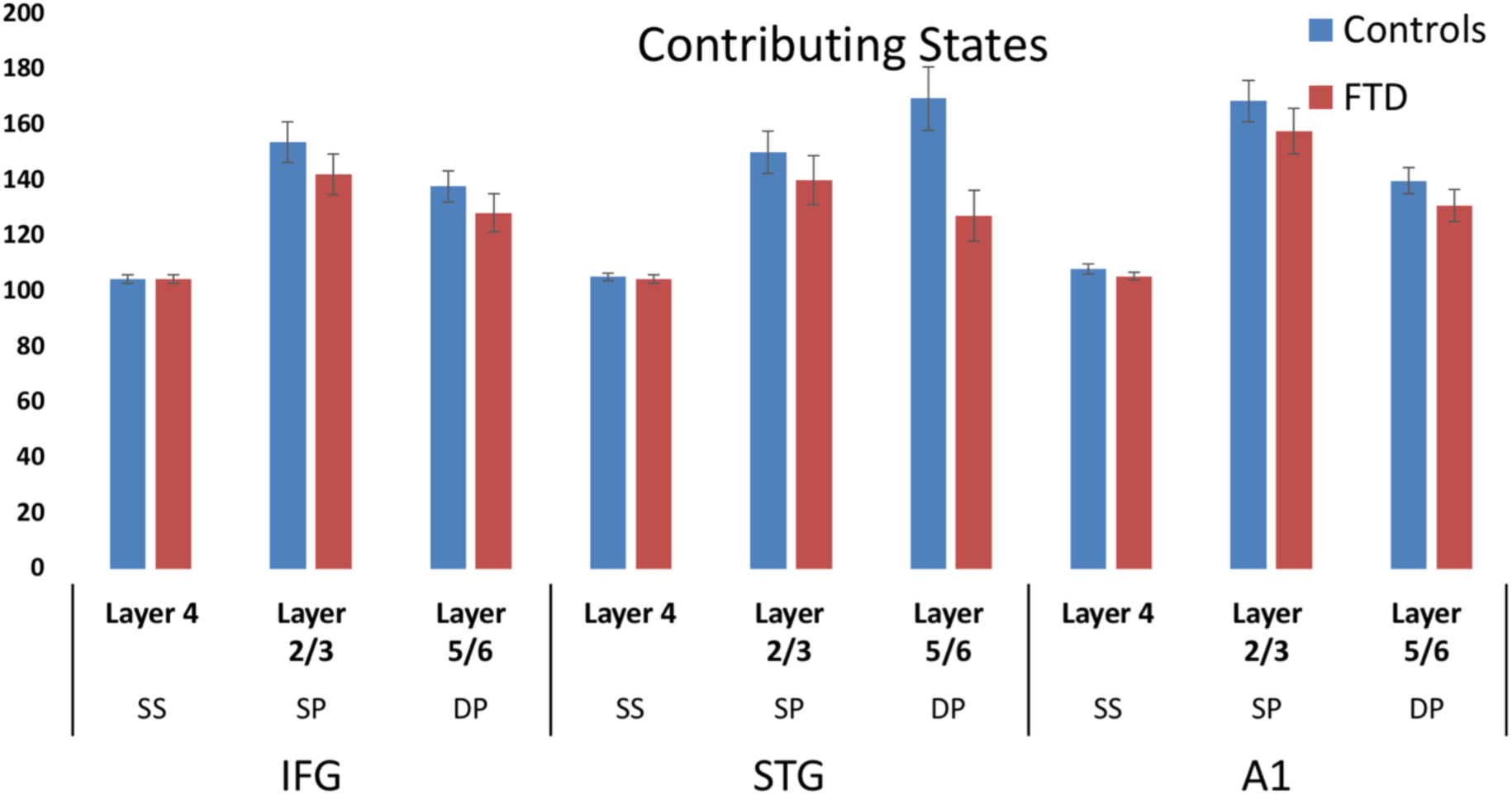
Right: Bar chart with error bars demonstrating the layer contributions per node (with enforced symmetry). Blue and red bars depict controls and bvFTD groups, respectively. Left: Scatter demonstrating layer 5/6 STG reduction in bvFTD compared with controls (red) and the trend in layer 2/3 IFG (blue).

**Figure 7.**
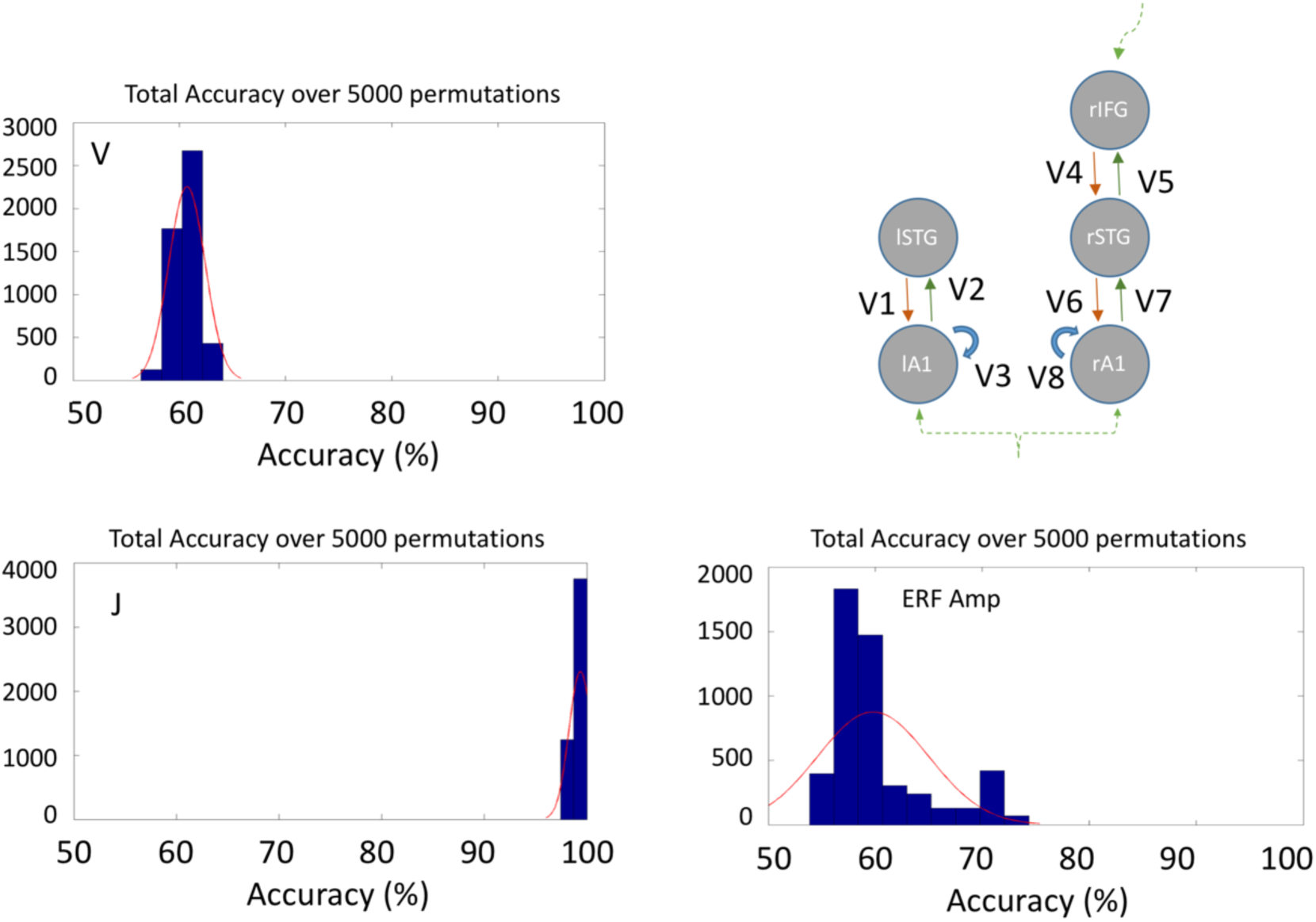
MMN-amplitude and model based classification. Histograms showing overall accuracy over 5000 permutations with leave-one-out. Note that for J the mean accuracy is 99.6%.

### Analysis B: Effective connectivity changes

Analysis of the posterior parameter estimates for intrinsic connectivity confirmed an increase in superficial layer (L2/3) pyramidal cell ‘inhibitory self gain’ (decay function) in the bvFTD group in the STG (p=.0257) along with a reduction in deep layer (L5/6) pyramidal cell self modulation in A1 (p=.0381) (Figure 8). Thus in effect superficial temporal regions exhibited hypoactive stimulus related activity while deep sensory regions exhibited a hyperactive sensory response.

**Figure 8.**
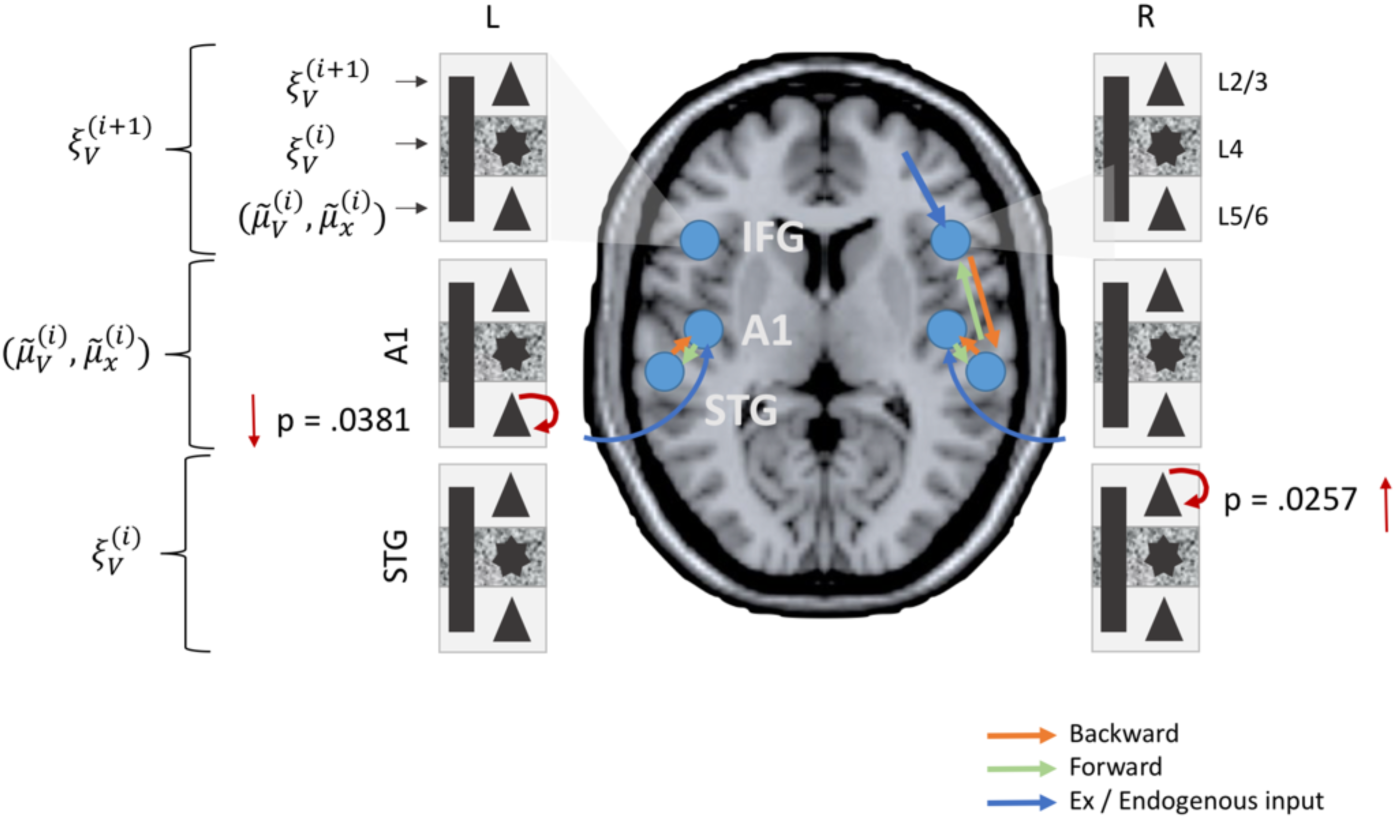
Local (intrinsic) parameter differences between bvFTD and controls. bvFTD show increases in L2/3 SP self-modulation in temporal areas (STG) and reductions in L5/6 SP self-modulation in sensory areas (A1).

## Discussion

This neurophysiological study of behavioural variant frontotemporal dementia has three principal results that contribute to an understanding of the disease. First, we replicate the observation that bvFTD reduces the amplitude of the mismatch negativity (Hughes and Rowe 2013), with patients failing to either adapt to predictable events and react to the unexpected events, compared with healthy adults. Second, we confirmed the neurophysiological prediction arising from the hypothesis of laminar selectivity of frontotemporal lobar degeneration (Kersaitis et al. 2004; Kim et al. 2012; Santillo and Englund 2014), in that bvFTD significantly reduces the contribution to the local electromagnetic signal from deep pyramidal cells (figure 6) and demonstrates a clear trend towards reduction in superficial layers, but not layer IV cells. Third, bvFTD causes faster decay of superficial layer pyramidal cells’ activity in superficial temporal areas and slower decay of deep-layer pyramidal cells in auditory cortex. We interpret these changes in terms of the way that sensory information is predicted in hierarchical frontotemporal networks: that the gain function of superficial pyramidal cells feeding prediction errors forward is reduced, with converse changes in the conditional expectations represented in lower level deep pyramidal populations.

The initial analysis of the event-related MMN replicates previous work in a smaller cohort (n=11)(Hughes and Rowe 2013). Such a global deficit in MMN generation is not unique to bvFTD, but has been reported in several neurological and psychiatric disorders (Mondragón-Maya et al. 2011; Naatanen et al. 2011). However, patients with bvFTD are unusual in the reduction of MMN to all deviant types tested, at the group level. However, the typical parameters used to describe the evoked MMN response (magnitude and latency) proved insufficient to enable accurate classification.

The model based approach taken using Dynamic Causal Modelling allows a richer parameterisation of the neurophysiologic response to standard and deviant tones, through generative networks in frontotemporal cortex. These parameters were optimised by inverting to the whole timeseries of the initial MMN (300ms), not merely the peak amplitude and latency. We built a moderately complex model that does not claim to include all regions in which a MMN is generated, but which includes six principal generators that have been most extensively studied by MEG, EEG and direct electrocorticography (Garrido et al. 2008, 2009). Critically, analysis of human MEG and electrocorticography confirms similar hierarchical network features. In this study however, we adopted the more complex and biologically informed canonical microcircuit model to examine the mechanism by which bvFTD alters the MMN.

With six principal regions in frontotemporal cortex, and possible modulation of feedforward and/or feedback connectivity by deviant versus standard stimuli, there are many possible models. We searched for the most likely model, from a principled set of 21 models, based on Phillips *et al*. (Phillips et al. 2015), which includes the model sub-set studied by Boly *et al*. (Yong Chen, Yuting Yang, Megan van Overbeek, Jill R. Donigian, Paul Baciu, Titia de Lange and Mammalian 2012) and Garrdio *et al*. (Garrido et al. 2008). We used hierarchical Bayesian model selection, with both Fixed- and Random-effects models. FFX and RFX models differ in the interpretation of their posterior probabilities, sensitivity to outlying subjects, and whether they accommodate heterogeneity in generative models among a cohort. In this study, FFX and RFX were in accord, revealing model 14 as the most likely. Garrido and colleagues (Garrido et al. 2008) previously demonstrated a closely related modelas best fit in a ‘roving’ mismatch paradigm in healthy subjects, but they did not test an identical model. As in the winning model here (model 14), Phillips *et al*. (Phillips et al. 2015), included models with top-down inputs to IFG, conveying high-level predictions or *expectation* of an event occurring, as opposed to low level predictions of stimulus features.

BMS was not performed separately for each group in this study, because potential group differences in the generative models are accommodated by the RFX approach. A whole-study BMS was performed in order to compare parameters of the generative and spatial models between groups, which requires that the groups have the same parameter set (and because model averaging leads to parameters with less biological interpretation). To use BMS to investigate differences in model architectures between groups would introduce the confound and ambiguity arising from group-by-model parameter differences. In the next sections, we discuss the insights arising from the group differences in the most likely network model.

Our primary hypothesis was that superficial and deep layers of the frontal cortex and temporal association cortex would show the largest reduction in their contribution to the regional electromagnetic signal. This prediction rests on the well-characterised pathology of bvFTD, in which layers 2 and 3 contain early pathogenic protein aggregates and cell loss in human and animal models (Kersaitis et al. 2004). Moreover, selective loss from layer 5 of Von Economo, fork- and surrounding pyramidal neurons occurs in bvFTD (Kim et al. 2012), with an estimated 70% reduction in cell number *post mortem* (Seeley 2008). This L5 atrophy is a hallmark of bvFTD pathology.

Our finding demonstrates this reduction *in vivo* in bvFTD patients, with two critical interpretations. First, in the context of bvFTD, neurophysiological evidence of L5 cell loss atrophy may be a biomarker specific to bvFTD, and preservation of layer 5 could be a priority for disease modifying treatments of bvFTD. Second, that the weighted forward-model linked to generative models of cortical networks can capture the characteristics of disease specific neurodegeneration, and this that might be upheld in other human dementias and neurodegenerative disorders, for which *in vivo* neurophysiological assays are necessarily indirect.

The generative canonical microcircuit model, in contrast to a spatial weighted forward-model, provides insights into the effect of bvFTD on intrinsic coupling connectivity within cortical regions. Two complementary changes were observed in bvFTD, compared with controls: (i) increased inhibitory auto-modulation of superficial layer pyramidal cells in STG, indicating a more rapid decay of activity in the absence of extrinsic driving inputs to the pyramidal cell population; and (ii) decreased auto-modulation of deep layer pyramidal cells in auditory cortex, indicating more stable firing rates of pyramidal cells here. These findings are particularly relevant because of the critical roles that these parameters have for predictive coding of events.

To understand the clinical consequences of these observations we interpret our findings within the predictive coding hypothesis (Rao and Ballard 1999; Friston 2005; Bastos et al. 2012), in which information about expectations (beliefs) and observed states (sensory inputs) are represented in a cortical hierarchy. Although the information content becomes more abstract and temporally extended in higher levels, the asymmetry between feedforward and feedback of information is analogous between hierarchical levels. Specifically, stellate cells in layer 4 receive feedforward connections that encode the prediction errors on the hidden causes of the level below. Superficial pyramidal cells encode and feed forward these prediction errors on hidden causes, whereas deep pyramidal cells encode the conditional expectations or belief, so as to elaborate feedback predictions to lower levels. Within our hierarchical model of bvFTD, the superficial temporal cortex are proposed to process changes in the physical properties of the tones in terms of the five variable dimensions of frequency, duration, amplitude, laterality, and temporal profile. In contrast, auditory cortex combines the predictions passed down from STG with the ‘raw’ sensory stream entering auditory layer 4.

The two parameter differences we see in the bvFTD group may therefore reflect one – single - integrated deficit; namely, a lack of *precision* in the encoding of prediction errors. This discrepancy in prediction subsequently propagates, leading to errors in the encoding of ‘conditional expectation’ in lower portions of the hierarchy (L5/6 encoding reduction in A1), which are observed macroscopically as a failure to generate a mismatch response.

We also tested whether the parameters of the generative model, in terms of intrinsic coupling within regions, or extrinsic coupling between regions, would provide a better biomarker of disease than the more typical summary features of the evoked mismatch response (amplitude and/or latency). This heuristic approach could be useful in determining whether model parameters offer robust biomarkers for stratification or outcome measures in future experimental medicine studies, using cohorts of a size and mixed pathology (Tau vs. TDP43) that is realistic for early phase trials.

The data clearly show that simple machine learning using a support vector machine provides highly accurate classification with model parameters of intrinsic coupling. This contrasts with the lower accuracy using MMN amplitude or intrinsic coupling parameters between regions. These latter methods supported above-chance classification, but the actual accuracy level (∼60%) would not be useful in a trials context, and suggests that these parameters are not sufficiently sensitive either as a diagnostic or prognostic biomarkerof bvFTD. The sensitivity and specificity of the Layer-by-node parameters in classification were 99.2 and 100%, respectively, making this a strong candidate marker. This finding has an added advantage over many imaging biomarkers in that the physical basis of the parameter is not merely an indirect correlate of the disease process, but rather reflects a component of the disease process itself – namely the reduction in the laminar output due to cell dysfunction and death.

The weaker classification accuracy using the between-region connectivity strengths (effective connectivity) was surprising in light of the findings of Brodersen *et al*. (Brodersen et al. 2011), who used a similar ‘generative embedding’ approach to distinguish between healthy and aphasic patients. However, they used a conceptually analogous but mechanistically distinct version of Dynamic Causal Modelling, for functional magnetic resonance imaging data. It is also possible that classification would have been higher if model selection was performed on each group separately, and subsequent models used for classification. However, such an approach is arguably biased towards a group difference in parameters, and we selected the model which best captured the pooled population rather than individual groups.

Future studies could extend our approach to include more biologically detailed generative models in experimental medicine studies and early phase trials. For example, a NMDA-receptor furnished conductance based model has been successfully used to model channelopathies in individual cases (Gilbert et al. 2016), and the effects of dopamine on working memory systems in the frontal cortex (Moran, Symmonds, et al. 2011). This would be especially relevant to the use assessment of target engagement of candidate therapies (Moran et al. 2013).

Dynamic Causal Models can in principle also incorporate pathological and structural anatomical information. For example, *post mortem* or selective PET-ligand data may separate cases with Tau pathology from TDP43 pathology, which are expected to be in roughly equal numbers in a bvFTD cohort. However, the current PET ligands lack demonstration of selectivity between Tau and TDP43 pathology, despite being sensitive to the burden and distribution of Tau pathology in FTD, progressive supranuclear palsy and Alzheimer’s disease (Bevan-Jones et al. 2017; Passamonti et al. 2017). The *post mortem* approach also requires time, to classify patients *post hoc*. From our cohort of 40 patients, 15 have died, and 5 underwent post mortem examination and three others have had genetic testing to indicate the molecular pathology.

Such models can also assess the generators of magneto- and electroencephalography signals at rest and in more complex task (Moran, Symmonds, et al. 2011), optimised by inverting to evoked responses as we did in this study, or the spectral density (Moran et al. 2009; Moran, Stephan, et al. 2011). However, the cognitive processes underlying variation at ‘rest’ are obscure, which confounds the interpretation of group differences in resting state data. Conversely, more complex tasks of social, economic, linguistic, mnemonic, affective or motor systems are of immediate relevance to the phenomenology of frontotemporal dementia (Hughes et al. 2011), but would require additional training and are subject to performance confounds. The MMN paradigm achieves a good compromise, of minimal set up and no training, while preserving a clear neurocognitive interpretation.

In conclusion, the inversion of generative models of cortical microcircuits, including laminar weighting of the spatial forward model to magnetoencephalography sensors, provides not only evidence of abnormal MMN responses in bvFTD, but also reveals two mechanisms by which the observed physiological response differs. Increasing the sophistication of human neurophysiological insights from MEG and EEG can provide heuristic biomarkers, but also facilitates cross-species comparisons between the physiology of transgenic models of frontotemporal lobar degeneration and their human disorders. We suggest that early phase clinical trials and experimental medicines studies consider integrating model based analysis of MEG and/or EEG, to understand the efficacy and mechanism of emerging candidate therapies.

## Acknowledgements.

This work was funded by the Wellcome Trust, the Medical Research Council (MC-A060-5PQ30), and the NIHR Cambridge Biomedical Research Centre. AS is supported by a Wellcome Trust Strategic Award (104943/Z/14/Z)

